# Cross-Platform Calibration of Epigenetic Age Between the EPICv2 and MSA Arrays

**DOI:** 10.64898/2025.12.10.693592

**Authors:** Yui Tomo, Tatsuma Shoji, Ryo Nakaki

## Abstract

Epigenetic clocks are regression models that predict biological age based on the DNA methylation patterns. They are developed using DNA methylation array data; however, systematic differences in the predicted values between the Infinium Methylation Screening Array (MSA) and Infinium MethylationEPIC v2.0 BeadChip (EPICv2) platforms have not been evaluated. We quantified the systematic differences between the MSA and EPICv2 arrays for six major epigenetic clocks (Horvath, Hannum, PhenoAge, GrimAge, GrimAge v2, and DunedinPACE) using 166 human blood samples measured on both platforms. For the five clocks expressed in years, the mean differences (MSA *−* EPICv2) ranged from *−*7.67 years for the Horvath clock to *−*0.634 years for GrimAge v2, despite strong correlations between platforms (*r* = 0.946–0.991). For DunedinPACE, which predicts the pace of aging, the mean difference was *−*0.117, with a correlation coefficient of 0.855 between the platforms. Based on these training data, we developed three linear regression models to map the MSA-based epigenetic ages and pace of aging to the EPICv2-based values, which were used as a reference: Model 1 (offset location calibration), Model 2 (slope and location calibration), and Model 3 (slope and location calibration with covariates). Validation using 48 independent samples measured at separate institutions showed that the calibration models reduced cross-platform differences for Horvath, Hannum, PhenoAge, GrimAge, and DunedinPACE. Although Model 3 yielded mean differences closest to zero for the Horvath clock (*−*0.227 years) and PhenoAge (*−*0.510 years), Model 1, the simplest specification, also reduced the mean cross-platform differences to *−*1.03 years for the Horvath clock and *−*1.37 years for PhenoAge. For DunedinPACE, Model 1 reduced the mean difference from *−*0.092 to 0.025 and the RMSE from 0.103 to 0.052. These methods are expected to enable reliable epigenetic age comparisons across platforms and facilitate large-scale epigenetic studies.

## 1. Introduction

DNA methylation is an epigenetic modification involving the covalent addition of methyl groups to cytosine bases in DNA molecules, particularly at cytosine-guanine dinucleotide (CpG) sites. These modifications are transmitted to daughter cells. DNA methylation regulates gene expression without altering the DNA sequence and is influenced by genetic, environmental, and aging factors [1, 2, 3, 4]. DNA methylation patterns are particularly susceptible to age-related changes. Analysis of 476, 366 CpG sites revealed that approximately 29% of these sites exhibited age-correlated methylation levels, with approximately 40% becoming hypermethylated and the remainder becoming hypomethylated with age [5]. Therefore, regression models that predict biological age based on DNA methylation patterns, known as epigenetic clocks, have been developed and play an important role in aging research [6, 7, 8].

Several epigenetic clocks have been developed to evaluate the various aspects of aging. The Horvath clock (2013), the first pan-tissue epigenetic clock applicable across multiple tissue types, was designed using 353 CpG sites to predict chronological age [6]. In contrast, the Hannum clock (2013) was developed for blood samples using 71 CpG sites to predict chronological age [9]. These first-generation clocks were designed to predict chronological age. Second-generation clocks have been developed to measure biological age, offering a better reflection of health status and mortality risk. These second-generation clocks include PhenoAge (2018), which predicts phenotypic age, incorporating blood biomarkers and mortality risk [10]; GrimAge (2019), a DNA methylation-based predictor of mortality risk, based on DNAm surrogates for smoking pack-years and plasma protein biomarkers, transformed to provide an output in years [11]; and GrimAge v2 (2022), which extends this framework by incorporating additional DNAm-based protein surrogates [12]. DunedinPACE (2022), a third-generation clock, provides a measure of the pace of aging [13].

DNA methylation arrays from Illumina Inc. (San Diego, CA, USA) are primarily used to measure methylation and develop epigenetic clocks. Illumina DNA methylation arrays have evolved from the Infinium Human-Methylation27 BeadChip (27K array) and Infinium HumanMethylation450 BeadChip (450K array) to the Infinium MethylationEPIC BeadChip (EPIC array), Infinium MethylationEPIC v2.0 BeadChip (EPICv2 array), and Infinium Methylation Screening Array (MSA array) [14, 15, 16]. The EPICv2 array covers over 930, 000 CpG sites, maintaining backward compatibility with the EPIC array while incorporating additional biologically relevant regions [17]. In contrast, the MSA array is a high-throughput platform that targets approximately 270, 000 CpG sites [16]. Although the MSA platform enables larger cohort studies and clinical applications, platform-specific technical characteristics may produce systematic differences in the methylation measurements when compared to other platforms.

Cross-platform differences can affect the reliability and reproducibility of epigenetic clocks. Different array platforms may yield varying methylation values for identical samples because of technical differences in probe design, chemistry, and the signal detection methods [18, 19]. These differences complicate the integrative analysis of data measured on different platforms and threaten the validity of longitudinal studies and meta-analyses [20]. Moreover, in clinical applications, cross-platform differences may lead to misestimation of individual biological ages, affecting disease risk predictions and intervention efficacy assessments. Previous studies have examined array-dependent differences among the 27K, 450K, EPICv1, and EPICv2 platforms, including analyses of Horvath and Hannum estimates across the 27K, 450K, and EPIC arrays [21], and more recent evaluations have focused on EPICv2 [22, 23, 24]. However, to our knowledge, a comparison of the epigenetic clock values measured by the EPICv2 and MSA arrays has not yet been reported.

This study aimed to quantify the platform differences between the EPICv2 and MSA arrays for six major epigenetic clocks (Horvath, Hannum, PhenoAge, GrimAge, GrimAge v2, and DunedinPACE). We measured 166 pairs of human blood samples on the two platforms and evaluated the differences in the values predicted by each epigenetic clock. Linear regression models were estimated to calibrate these differences in the epigenetic ages calculated for the MSA array. The calibration methods were validated using 48 independent samples measured at different institutions. The results are expected to help to improve the compatibility of epigenetic age data obtained from different platforms and enhance the reliability of longitudinal studies and meta-analyses.

The remainder of this paper is organized as follows: In Section 2, we describe the procedures for sample collection and DNA methylation analysis and the methods used to evaluate and calibrate cross-platform differences in epigenetic clock values. In Section 3, we present the cross-platform differences before and after calibration and the estimated calibration models. In Section 4, we discuss the utility and limitations of the calibration methods and directions for future research.

## 2. Materials and Methods

### 2.1. Sample Collection and Preparation

Whole-blood samples from healthy adult volunteers in Japan were obtained through two independently approved observational studies. In total, 214 unique individuals were included in this study. Two non-overlapping datasets were generated: a training dataset with 166 individuals and a validation dataset with 48 individuals. The 166 blood samples in the training dataset were collected at Y’s Science Clinic Hiroo (Minato-ku, Tokyo, Japan) as part of the research project “Evaluation of Biological Age Based on DNA Methylation and Its Clinical Significance.” This study was reviewed and approved by the Shiba Palace Clinic Institutional Review Board (approval number 155728_rn-37828, approval on 11, July, 2024), and written informed consent was obtained from all the participants. The 48 blood samples in the validation dataset were collected as part of the independent clinical research project “Evaluation of Epigenetic Clock Measurement in Clinical Practice.” This study was reviewed and approved by the Asia-Oceania AntiAging Promotion Association Ethics Review Committee (approval number AOAAPA22-001, approved on May 27, 2022). Written informed consent was obtained from all participants. Genomic DNA was extracted using the Maxwell RSC Blood DNA Kit (Promega, Madison, WI, USA) according to the manufacturer’s protocol. DNA concentrations were assessed using a NanoDrop spectrophotometer (Thermo Fisher Scientific, Minato-ku, Tokyo, Japan), and samples were stored at *−*80*^◦^*C until bisulfite conversion. All procedures adhered to the Ethical Guidelines for Medical and Biological Research Involving Human Subjects, issued by the Government of Japan.

### 2.2. DNA Methylation Analysis

Genome-wide DNA methylation profiling was performed using two Illumina BeadChip platforms: EPICv2 and MSA. For the training dataset (166 samples), EPICv2 measurements were generated by Macrogen Inc. (Gangnamgu, Seoul, Republic of Korea). MSA measurements were performed at Rhelixa Inc. (Chuo-ku, Tokyo, Japan) using aliquots derived from the same extracted DNA sample. For the validation dataset (48 samples), EPICv2 measurements were generated by Novogene Co., Ltd. (Chaoyang District, Beijing, China), and MSA measurements were obtained from CD Genomics (Shirley, NY, USA). The two platforms were independently processed. No individual appeared in both datasets. Bisulfite conversion was conducted using the EZ DNA Methylation Kit (Zymo Research, Irvine, CA, USA) according to the manufacturer’s protocol. The converted DNA was amplified, fragmented, and hybridized to each array, after which the arrays were scanned using an Illumina iScan System.

Raw IDAT files were processed using the SeSAMe pipeline. Background calibration and probe-level masking were performed using the “QCDPB” pipeline in SeSAMe. The pipeline included normal-exponential out-of-band (noob) background calibration and the pOOBAH detection mask, with the pOOBAH p-value threshold set to 0.05 [25, 26]. Beta-values were calculated as the ratio of methylated signal intensity to total signal intensity. Probe suffixes were collapsed to the probe-prefix level by averaging, and beta-values masked by the quality control (QC) were retained as missing values. Rows corresponding to control and background probes, identified by probe IDs beginning with “ctl” or “cgBK,” were removed from the final beta matrix.

Epigenetic clocks (Horvath, Hannum, PhenoAge, GrimAge, GrimAge v2, and DunedinPACE) were computed separately for EPICv2 and MSA using the processed beta-values. Missing values were handled using standard methods implemented in Biolearn. Specifically, for Horvath, Hannum, PhenoAge, GrimAge, and GrimAge v2, the sesame 450k gold standard values were used to impute missing values, whereas for DunedinPACE, the DunedinPACE gold standard values were used [27]. No normalization to an external gold-standard reference dataset was applied before clock calculation for the other five clocks. For each of the epigenetic clocks and arrays, the number of missing CpGs before QC was calculated (required CpGs *−* CpGs on the array). Furthermore, the number of CpGs affected by QC masking in at least one sample was counted for each epigenetic clock and dataset. For DunedinPACE, the probe coverage and number of CpGs affected by QC masking were evaluated separately for the 173 model CpGs and the full 20,000-CpG preprocessing set.

The software versions were as follows: R 4.5.3, SeSAMe 1.28.1, sesameData 1.28.0, BiocParallel 1.44.0, preprocessCore 1.72.0, Python 3.11.15, and biolearn 0.9.1. The computational environment was fixed using Docker, and the Dockerfile and the full pipeline scripts are available at https://github.com/tatsumashoji/methylation-beta-pipeline/tree/paper_msa_epicv2_grimagev1.

### 2.3. Evaluation of Cross-Platform Differences

Systematic differences between platforms were evaluated using the 166 training samples. For each epigenetic clock, the difference between the MSA and EPICv2 array-based values was computed. Scatter and Bland–Altman plots were generated for visual evaluation.

### 2.4. Development of Calibration Models

We used linear regression models to develop calibration methods to map the epigenetic ages and pace of aging calculated from MSA to those calculated from EPICv2. The EPICv2-based values were treated as a reference for cross-platform calibration, not as a gold-standard measure of the underlying biological age. For each epigenetic clock, EPICv2-based values were regressed on MSA-based values using the 166 training samples. We used three model specifications: Let *Y*_EPICv2_ *∈* R*_≥_*_0_ denote the EPICv2-based clock value, *Y*_MSA_ *∈* R*_≥_*_0_ denote the MSA-based clock value, *X*_Age_ *∈* R*_≥_*_0_ denote the chronological age, and *X*_Sex_ *∈ {*0, 1*}* denote a binary variable for sex. Sex was coded as 0 for female and 1 for male. Let *β*_0_*, β*_1_*, β*_2_*, β*_3_ *∈* R denote the regression coefficient parameters. The mean structures of the three specified linear regression models are expressed as follows:

(i) Model 1. Offset location calibration:

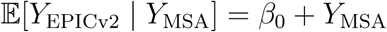

(ii) Model 2. Slope and location calibration:

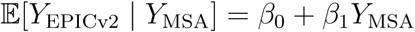

(iii) Model 3. Slope and location calibration with covariates:

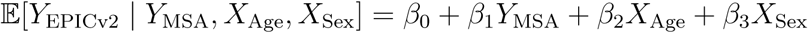

Regression coefficient parameters were estimated using ordinary least squares, and estimates were reported using the standard errors, t-statistics, and p-values from the t-tests.

The developed regression models were evaluated on the validation datasets comprising 48 samples collected from different institutions. The mean absolute error (MAE), root mean squared error (RMSE), and difference of the calibrated and uncalibrated MSA-based values in the validation dataset were determined relative to the EPICv2-based clock values, and the correlation coefficients were calculated. Scatter and Bland–Altman plots were generated.

## 3. Results

### 3.1. Sample Characteristics

The training dataset consisted of 166 blood samples from healthy adults, including 92 males (55.4%) and 74 females (44.6%) (Table 1). The mean chronological age was 47.1 years (standard deviation [SD]: 12.3 years; range 24.6–76.8 years). The validation dataset consisted of an independent set of 48 participants, including 24 males (50.0%) and 24 females (50.0%) (Table 2). The mean chronological age was 50.5 years (SD: 14.3 years; range 23.6–74.5 years).

**Table 1:** Baseline characteristics of the training dataset (*n* = 166).

| Characteristic | Value |  |
| --- | --- | --- |
| **Participant characteristics** |  |  |
| Chronological age (years) | 47.1 (12.3) |  |
| Chronological age range (years) | 24.6–76.8 |  |
| Male, *n* (%) | 92 (55.4%) |  |
| Female, *n* (%) | 74 (44.6%) |  |
| **Epigenetic age, mean (SD)** |  |  |
|  | EPICv2 | MSA |
| Horvath | 54.45 (11.57) | 46.78 (12.12) |
| Hannum | 48.57 (11.30) | 44.21 (11.39) |
| PhenoAge | 40.74 (13.29) | 35.65 (13.29) |
| GrimAge | 47.93 (10.17) | 46.63 (10.05) |
| GrimAge v2 | 53.30 (10.08) | 52.66 (10.13) |
| DunedinPACE | 1.02 (0.09) | 0.90 (0.10) |

**Table 2:** Baseline characteristics of the validation dataset (*n* = 48).

| Characteristic | Value |  |
| --- | --- | --- |
| **Participant characteristics** |  |  |
| Chronological age (years) | 50.5 (14.3) |  |
| Chronological age range (years) | 23.6–74.5 |  |
| Male, *n* (%) | 24 (50.0%) |  |
| Female, *n* (%) | 24 (50.0%) |  |
| **Epigenetic age, mean (SD)** |  |  |
|  | EPICv2 | MSA |
| Horvath | 56.21 (12.95) | 47.51 (13.17) |
| Hannum | 52.45 (12.17) | 48.53 (12.31) |
| PhenoAge | 43.22 (14.12) | 36.76 (14.40) |
| GrimAge | 52.43 (11.51) | 50.76 (11.84) |
| GrimAge v2 | 56.53 (11.24) | 56.56 (11.67) |
| DunedinPACE | 1.01 (0.09) | 0.92 (0.10) |

The number of CpGs missing before QC for each clock and platform is shown in Table 3. Coverage was calculated relative to the number of CpGs required by each clock. The number of clock CpGs affected by QC masking in at least one sample is shown in Table 4. For DunedinPACE, the 173 model CpGs and the full 20,000-CpG preprocessing set were reported separately.

**Table 3:** Coverage of CpGs required by each clock before quality control.

| Clock | Required | EPICv2 |  | MSA |  |
| --- | --- | --- | --- | --- | --- |
|  |  | Present | Missing | Present | Missing |
| Horvath | 353 | 340 (96.3%) | 13 (3.7%) | 347 (98.3%) | 6 (1.7%) |
| Hannum | 71 | 64 (90.1%) | 7 (9.9%) | 68 (95.8%) | 3 (4.2%) |
| PhenoAge | 513 | 495 (96.5%) | 18 (3.5%) | 510 (99.4%) | 3 (0.6%) |
| GrimAge | 1,030 | 845 (82.0%) | 185 (18.0%) | 1,010 (98.1%) | 20 (1.9%) |
| GrimAge v2 | 1,030 | 845 (82.0%) | 185 (18.0%) | 1,010 (98.1%) | 20 (1.9%) |
| DunedinPACE | 173 | 144 (83.2%) | 29 (16.8%) | 171 (98.8%) | 2 (1.2%) |
| (model CpGs) |  |  |  |  |  |
| DunedinPACE | 20,000 | 17,387 (86.9%) | 2,613 (13.1%) | 9,678 (48.4%) | 10,322 (51.6%) |
| (preprocessing set) |  |  |  |  |  |

**Table 4:** Clock CpGs affected by quality-control masking in at least one sample for the training (*n* = 166) and validation (*n* = 48) datasets.

| Clock | EPICv2 |  | MSA |  |
| --- | --- | --- | --- | --- |
|  | Training | Validation | Training | Validation |
| Horvath | 1 (0.3%) | 5 (1.4%) | 3 (0.8%) | 5 (1.4%) |
| Hannum | 0 (0.0%) | 1 (1.4%) | 0 (0.0%) | 0 (0.0%) |
| PhenoAge | 2 (0.4%) | 11 (2.1%) | 7 (1.4%) | 4 (0.8%) |
| GrimAge | 23 (2.2%) | 44 (4.3%) | 54 (5.2%) | 53 (5.1%) |
| GrimAge v2 | 23 (2.2%) | 44 (4.3%) | 54 (5.2%) | 53 (5.1%) |
| DunedinPACE | 2 (1.2%) | 5 (2.9%) | 3 (1.7%) | 2 (1.2%) |
| (model CpGs) |  |  |  |  |
| DunedinPACE | 2 (< 0.1%) | 185 (0.9%) | 3 (< 0.1%) | 261 (1.3%) |
| (preprocessing set) |  |  |  |  |

### 3.2. Evaluation of Cross-Platform Differences

A cross-platform comparison revealed clear systematic differences between the MSA and EPICv2 arrays for Horvath, Hannum, PhenoAge, and GrimAge, whereas the differences for GrimAge v2 were comparatively small. The mean difference between the MSA and EPICv2 arrays (MSA *−* EPICv2) for each clock in the training dataset (*n* = 166) is summarized in Table 5. The mean differences were *−*7.67 years for the Horvath clock and *−*5.09 years for PhenoAge, indicating that MSA yielded systematically lower values for these clocks. For GrimAge and GrimAge v2, the mean differences were *−*1.30 and *−*0.634 years, respectively, indicating comparatively small differences. For DunedinPACE, the mean difference was *−*0.117, again indicating that MSA produced lower values than EPICv2.

**Table 5:** Systematic differences (MSA *−* EPICv2) between the MSA and EPICv2 arrays.

| Epigenetic clock | Mean difference | SD |
| --- | --- | --- |
| Horvath | -7.67 | 3.43 |
| Hannum | -4.36 | 1.90 |
| PhenoAge | -5.09 | 4.38 |
| GrimAge | -1.30 | 1.38 |
| GrimAge v2 | -0.634 | 1.41 |
| DunedinPACE | -0.117 | 0.052 |

The scatter plots in Figure 1 show strong correlations between EPICv2 and MSA for all clocks (correlation coefficients 0.855 to 0.991), while revealing the presence of systematic offsets. In the Bland–Altman plots shown in Figure 1, most clocks exhibited approximately constant difference patterns that did not depend on the magnitude of the data.

**Figure 1:**
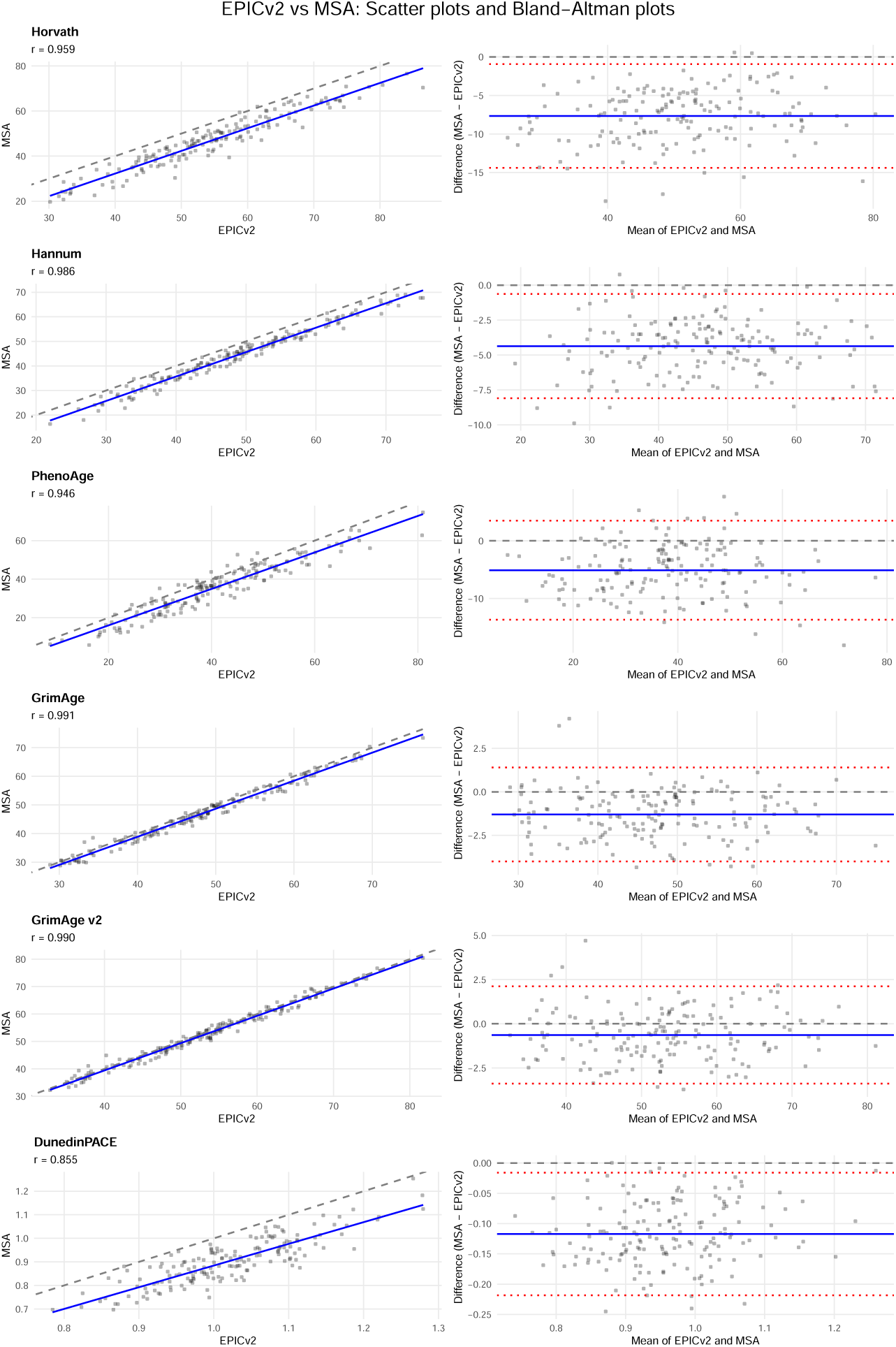
Comparison of epigenetic ages between the EPICv2 and MSA arrays on the training dataset (*n* = 166). Left column: scatter plots showing the relationship between EPICv2 (*x*-axis) and MSA (*y*-axis) values for each epigenetic clock. The dashed line represents the identity line (*y* = *x*), and the blue line shows the linear regression fit. Right column: Bland–Altman plots displaying the difference between MSA and EPICv2 values (*y*-axis) against the means (*x*-axis). The solid blue line indicates the mean difference, the dashed line represents zero difference, and the dotted red lines show the 95% limits of agreement (mean *±* 1.96SD). Each row represents a different epigenetic clock: Horvath, Hannum, PhenoAge, GrimAge, GrimAge v2, and DunedinPACE.

### 3.3. Development of Calibration Models

Three calibration models were developed using the training dataset (*n* = 166). The estimated regression coefficients are presented in Table 6. Model 1, the simplest model, added a constant term to the MSA array values. For example, the estimated intercepts were 7.67 years for the Horvath clock and 4.36 years for the Hannum clock. Model 2 included the MSA-based clock value as the sole predictor and estimated both the intercept and slope. For all clocks, the estimated regression coefficient for *Y*_MSA_ ranged from 0.793 (DunedinPACE) to 1.00 (GrimAge). Model 3 included chronological age and sex as additional explanatory variables, in addition to the MSA array value. The estimated chronological-age coefficients were positive for all clocks and were 0.339 for the Horvath clock and 0.366 for PhenoAge. The estimated sex coefficients were 1.30 for the Horvath clock and 0.968 for the Hannum clock.

**Table 6:**
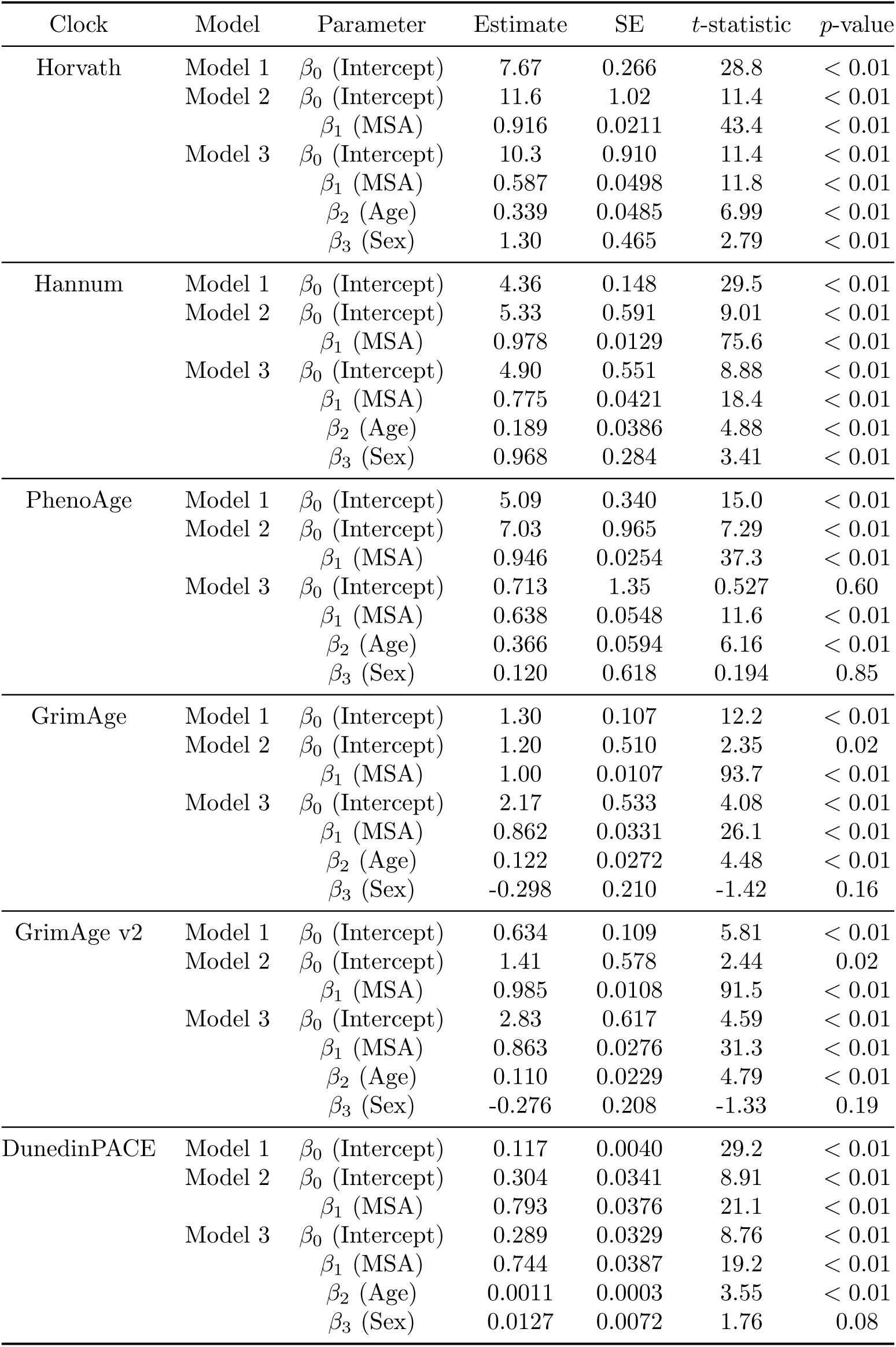
Regression coefficients of the calibration models.

| Clock | Model | Parameter | Estimate | SE | *t*-statistic | *p*-value |
| --- | --- | --- | --- | --- | --- | --- |
| Horvath | Model 1 | $\beta_0$ (Intercept) | 7.67 | 0.266 | 28.8 | < 0.01 |
| | Model 2 | $\beta_0$ (Intercept) | 11.6 | 1.02 | 11.4 | < 0.01 |
| | | $\beta_1$ (MSA) | 0.916 | 0.0211 | 43.4 | < 0.01 |
| | Model 3 | $\beta_0$ (Intercept) | 10.3 | 0.910 | 11.4 | < 0.01 |
| | | $\beta_1$ (MSA) | 0.587 | 0.0498 | 11.8 | < 0.01 |
| | | $\beta_2$ (Age) | 0.339 | 0.0485 | 6.99 | < 0.01 |
| | | $\beta_3$ (Sex) | 1.30 | 0.465 | 2.79 | < 0.01 |
| Hannum | Model 1 | $\beta_0$ (Intercept) | 4.36 | 0.148 | 29.5 | < 0.01 |
| | Model 2 | $\beta_0$ (Intercept) | 5.33 | 0.591 | 9.01 | < 0.01 |
| | | $\beta_1$ (MSA) | 0.978 | 0.0129 | 75.6 | < 0.01 |
| | Model 3 | $\beta_0$ (Intercept) | 4.90 | 0.551 | 8.88 | < 0.01 |
| | | $\beta_1$ (MSA) | 0.775 | 0.0421 | 18.4 | < 0.01 |
| | | $\beta_2$ (Age) | 0.189 | 0.0386 | 4.88 | < 0.01 |
| | | $\beta_3$ (Sex) | 0.968 | 0.284 | 3.41 | < 0.01 |
| PhenoAge | Model 1 | $\beta_0$ (Intercept) | 5.09 | 0.340 | 15.0 | < 0.01 |
| | Model 2 | $\beta_0$ (Intercept) | 7.03 | 0.965 | 7.29 | < 0.01 |
| | | $\beta_1$ (MSA) | 0.946 | 0.0254 | 37.3 | < 0.01 |
| | Model 3 | $\beta_0$ (Intercept) | 0.713 | 1.35 | 0.527 | 0.60 |
| | | $\beta_1$ (MSA) | 0.638 | 0.0548 | 11.6 | < 0.01 |
| | | $\beta_2$ (Age) | 0.366 | 0.0594 | 6.16 | < 0.01 |
| | | $\beta_3$ (Sex) | 0.120 | 0.618 | 0.194 | 0.85 |
| GrimAge | Model 1 | $\beta_0$ (Intercept) | 1.30 | 0.107 | 12.2 | < 0.01 |
| | Model 2 | $\beta_0$ (Intercept) | 1.20 | 0.510 | 2.35 | 0.02 |
| | | $\beta_1$ (MSA) | 1.00 | 0.0107 | 93.7 | < 0.01 |
| | Model 3 | $\beta_0$ (Intercept) | 2.17 | 0.533 | 4.08 | < 0.01 |
| | | $\beta_1$ (MSA) | 0.862 | 0.0331 | 26.1 | < 0.01 |
| | | $\beta_2$ (Age) | 0.122 | 0.0272 | 4.48 | < 0.01 |
| | | $\beta_3$ (Sex) | -0.298 | 0.210 | -1.42 | 0.16 |
| GrimAge v2 | Model 1 | $\beta_0$ (Intercept) | 0.634 | 0.109 | 5.81 | < 0.01 |
| | Model 2 | $\beta_0$ (Intercept) | 1.41 | 0.578 | 2.44 | 0.02 |
| | | $\beta_1$ (MSA) | 0.985 | 0.0108 | 91.5 | < 0.01 |
| | Model 3 | $\beta_0$ (Intercept) | 2.83 | 0.617 | 4.59 | < 0.01 |
| | | $\beta_1$ (MSA) | 0.863 | 0.0276 | 31.3 | < 0.01 |
| | | $\beta_2$ (Age) | 0.110 | 0.0229 | 4.79 | < 0.01 |
| | | $\beta_3$ (Sex) | -0.276 | 0.208 | -1.33 | 0.19 |
| DunedinPACE | Model 1 | $\beta_0$ (Intercept) | 0.117 | 0.0040 | 29.2 | < 0.01 |
| | Model 2 | $\beta_0$ (Intercept) | 0.304 | 0.0341 | 8.91 | < 0.01 |
| | | $\beta_1$ (MSA) | 0.793 | 0.0376 | 21.1 | < 0.01 |
| | Model 3 | $\beta_0$ (Intercept) | 0.289 | 0.0329 | 8.76 | < 0.01 |
| | | $\beta_1$ (MSA) | 0.744 | 0.0387 | 19.2 | < 0.01 |
| | | $\beta_2$ (Age) | 0.0011 | 0.0003 | 3.55 | < 0.01 |
| | | $\beta_3$ (Sex) | 0.0127 | 0.0072 | 1.76 | 0.08 |

The performance of the calibration models in the validation dataset (*n* = 48) is summarized in Table 7. Model 1 calibrations reduced the differences for Horvath, Hannum, PhenoAge, GrimAge, and DunedinPACE, but not for GrimAge v2, which already showed a small difference in the validation dataset. The difference between the Horvath clock values was largely reduced from *−*8.70 to *−*1.03 years, and the MAE decreased from 8.70 to 2.49. The difference between the PhenoAge values was reduced from *−*6.46 to *−*1.37 years. For Model 2, the smallest differences were obtained for the Hannum clock (0.349 years). Model 3, which incorporated covariates, further improved the predictive performance. The MAE and RMSE reached their lowest values for many clocks, with notable improvements for PhenoAge (MAE: 3.58; RMSE: 4.36). The correlation coefficients also improved slightly, particularly for Horvath (0.977), PhenoAge (0.952), GrimAge (0.990), and GrimAge v2 (0.990).

**Table 7:** Model performance in the validation dataset. The mean difference was defined as the calibrated MSA-based value *−* the EPICv2-based value. MAE: mean absolute error; RMSE: root mean squared error; *r*: correlation coefficient.

| Clock | Model | MAE | RMSE | Mean Difference | $r$ |
| --- | --- | --- | --- | --- | --- |
| Horvath | Unadjusted | 8.70 | 9.21 | -8.70 | 0.973 |
|  | Model 1 | 2.49 | 3.19 | -1.03 | 0.973 |
|  | Model 2 | 2.62 | 3.21 | -1.09 | 0.973 |
|  | Model 3 | 2.20 | 2.72 | -0.227 | 0.977 |
| Hannum | Unadjusted | 3.92 | 4.18 | -3.92 | 0.993 |
|  | Model 1 | 1.23 | 1.52 | 0.443 | 0.993 |
|  | Model 2 | 1.21 | 1.47 | 0.349 | 0.993 |
|  | Model 3 | 1.14 | 1.44 | 0.072 | 0.993 |
| PhenoAge | Unadjusted | 6.78 | 8.42 | -6.46 | 0.927 |
|  | Model 1 | 4.40 | 5.57 | -1.37 | 0.927 |
|  | Model 2 | 4.35 | 5.45 | -1.44 | 0.927 |
|  | Model 3 | 3.58 | 4.36 | -0.510 | 0.952 |
| GrimAge | Unadjusted | 2.11 | 2.55 | -1.67 | 0.986 |
|  | Model 1 | 1.53 | 1.96 | -0.366 | 0.986 |
|  | Model 2 | 1.54 | 1.97 | -0.357 | 0.986 |
|  | Model 3 | 1.38 | 1.76 | -0.498 | 0.990 |
| GrimAge v2 | Unadjusted | 1.50 | 1.87 | 0.036 | 0.987 |
|  | Model 1 | 1.59 | 1.98 | 0.670 | 0.987 |
|  | Model 2 | 1.54 | 1.92 | 0.613 | 0.987 |
|  | Model 3 | 1.38 | 1.69 | 0.532 | 0.990 |
| DunedinPACE | Unadjusted | 0.093 | 0.103 | -0.092 | 0.875 |
|  | Model 1 | 0.041 | 0.052 | 0.025 | 0.875 |
|  | Model 2 | 0.040 | 0.049 | 0.021 | 0.875 |
|  | Model 3 | 0.040 | 0.051 | 0.024 | 0.862 |

For all calibration models and clocks, the calibrated MSA values were clustered along the identity line (Figure 2). For all models, the difference was centered around zero after calibration, as shown in the Bland–Altman plots in Figure 3.

**Figure 2:**
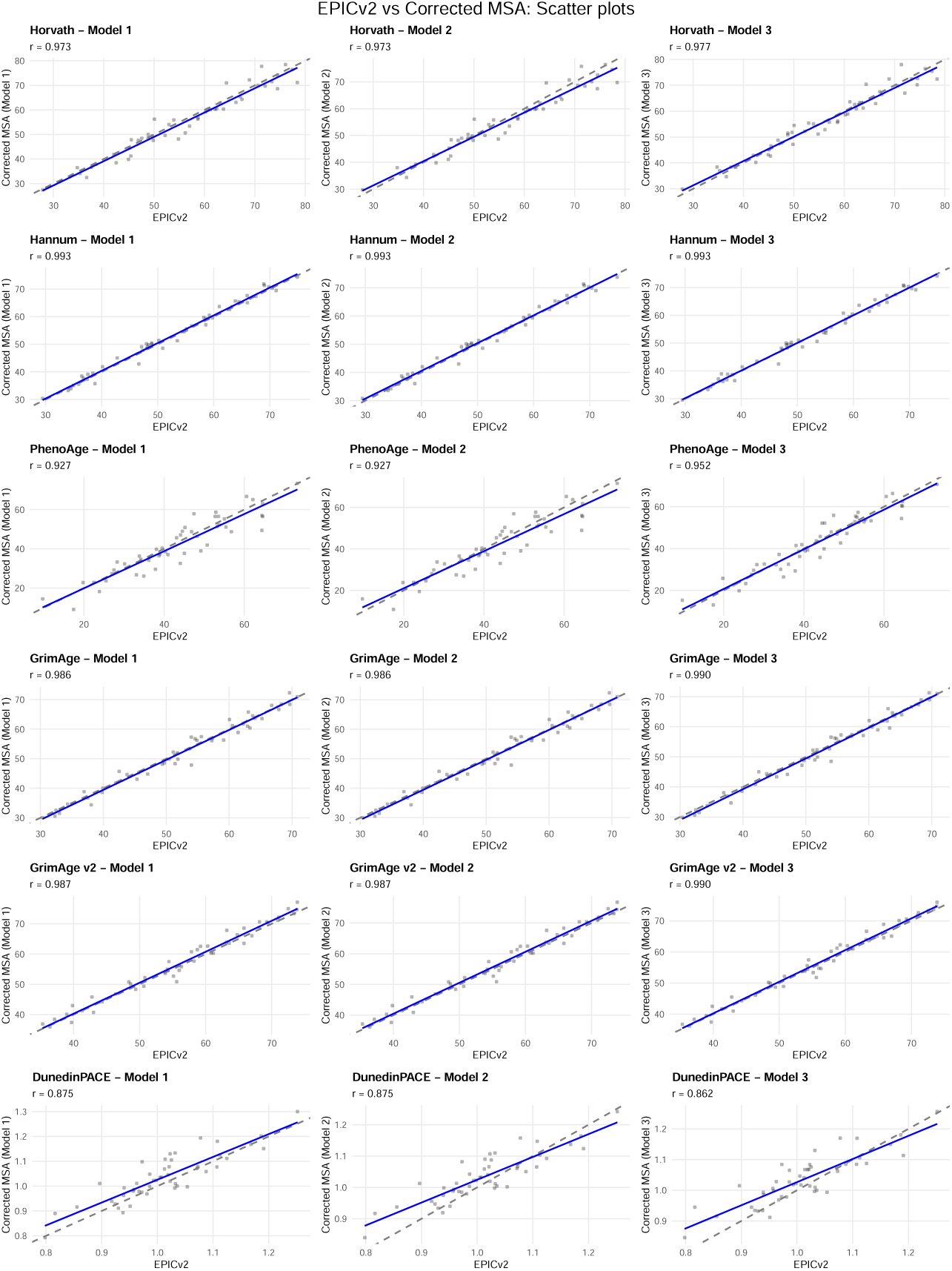
Scatter plots comparing EPICv2 values with calibrated MSA values after applying three calibration models to the validation dataset (*n* = 48). Each column represents a different calibration model: Model 1 (offset location calibration), Model 2 (slope and location calibration), and Model 3 (slope and location calibration with covariates). Each row corresponds to a different epigenetic clock. The dashed line represents the identity line (*y* = *x*), and the blue line shows the linear regression fit. The correlation coefficient (*r*) is displayed for each plot.

**Figure 3:**
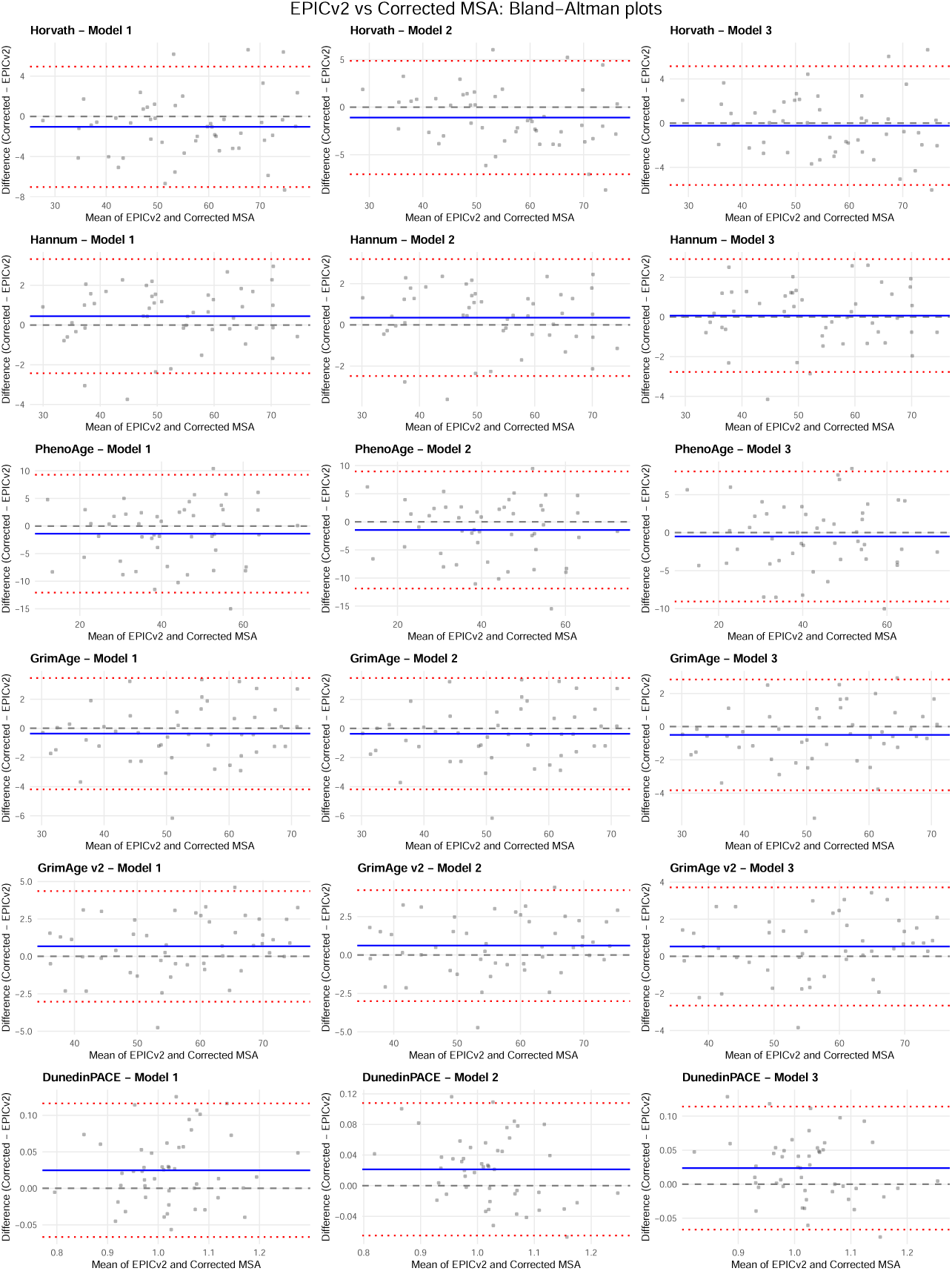
Bland–Altman plots showing the error structure after applying calibration models to MSA values, on the validation dataset (*n* = 48). Each column represents a different calibration model: Model 1 (offset location calibration), Model 2 (slope and location calibration), and Model 3 (slope and location calibration with covariates). Each row corresponds to a different epigenetic clock. The *y*-axis shows the difference between the calibrated MSA and EPICv2 values, and the *x*-axis shows their mean. The solid blue line indicates the mean difference, the dashed line represents zero difference, and the dotted red lines show the 95% limits of agreement.

## 4. Discussion

In this study, we quantified the systematic differences between the EPICv2 and MSA arrays across six major epigenetic clocks and developed and validated calibration methods to address these differences. Non-negligible systematic differences were identified between the platforms for Horvath, Hannum, PhenoAge, GrimAge, and DunedinPACE. The mean differences were *−*7.67 years for the Horvath clock and *−*0.117 for DunedinPACE. These findings demonstrate the difficulty of directly comparing epigenetic ages measured on different platforms and emphasize the need for calibration. Importantly, EPICv2 was used here as a reference platform for calibration rather than as a gold standard for the “true” biological age, and the results should be interpreted accordingly. Our objective was to quantify and correct the discrepancies in the epigenetic ages and pace of aging between the EPICv2 and MSA platforms to facilitate comparability across these platforms.

The calibration models effectively reduced the systematic differences for clocks with clear cross-platform differences, particularly the Horvath, Hannum, PhenoAge, GrimAge, and DunedinPACE clocks. Model 3, which incorporated slope and location calibrations with covariates, tended to perform best when chronological age and sex were included as regression covariates. For PhenoAge and GrimAge, Model 3 improved the MAE and RMSE and slightly improved the correlation coefficients. For DunedinPACE, Model 2 yielded the smallest RMSE, although Model 1 showed similar performance. This suggests that platform-related differences in epigenetic age may be associated with simple linear relationships and biological factors, such as age and sex. However, the differences in performance among the three models were relatively small, and even Model 1 (constant offset calibration), which was the simplest model, provided sufficient calibration. For example, applying Model 1 to the PhenoAge values largely reduced the difference from *−*6.46 to *−*1.37 years and reduced the mean absolute error from 6.78 to 4.40.

This demonstrates that sufficient accuracy can be achieved using a simple calibration method that provides a constant offset. The direction and magnitude of the cross-platform differences varied among the clocks. While the Horvath, Hannum, and PhenoAge clocks showed systematically lower values for MSA than for EPICv2, GrimAge showed a smaller negative difference, and GrimAge v2 exhibited a difference close to zero, indicating relatively good cross-platform comparability. These differences are likely attributable to the variables and CpG sites used by each epigenetic clock and the differences in the technical characteristics of the probes for these sites.

The relatively small differences observed for GrimAge and GrimAge v2 can be partially explained by their construction. Both models were trained by fitting a Cox proportional hazards model to time-to-death data using methylation-based surrogate markers for plasma proteins and smoking pack-years, with chronological age and sex included as predictors. The resulting linear predictor was then rescaled to the units of chronological age [11, 12]. Because chronological age is directly involved in the linear predictor, the relative contribution of the methylation-based components to the predicted value may be reduced.

Epigenetic clocks have been applied in various fields, including aging research, disease risk assessment, and the evaluation of intervention effects [28, 29, 30, 31]. The correction methods developed in this study may provide a practical approach for improving the comparability of MSA-based clock values with EPICv2-referenced values.

Epigenetic age acceleration (EAA), typically defined as the residual from the regression of epigenetic age on chronological age within a given dataset, is widely used as a predictor or outcome in association analysis [28, 32]. Residualization absorbs additive shifts, so correcting for a constant cross-platform offset is not necessary for analyses based on EAA. Nevertheless, calibration on the absolute age scale remains necessary for several reasons.

First, EAA is a relative quantity; the same epigenetic age measurement in the same individual yields different EAA values depending on the reference population used to define the residuals, so EAA values cannot be directly compared across cohorts or across platform combinations in meta-analyses of mixed platforms. Second, in longitudinal and interventional studies where an effect is interpreted as being “equivalent to some years of aging,” the results can be reported more directly using the absolute age scale, rather than the residual scale. Third, in settings such as forensic age estimation where chronological age is not available [33, 34], the predicted age must be reported on the year scale. The direct calibration of the epigenetic age is therefore necessary in these situations.

We considered the principal component (PC) versions of the epigenetic clocks proposed by Higgins-Chen et al. [35]. These PC clocks improve reliability by projecting CpG-level methylation data onto principal components estimated from a training dataset and then applying the clock model to the resulting PC scores. However, the MSA array does not contain the full set of CpGs required to reconstruct the predefined PC scores directly. Applying PC clocks to MSA data would therefore require additional imputation to reconstruct the predefined PC scores. Alternatively, PC clocks could be retrained for the MSA array, but this would require a large training dataset and would define a different model. Therefore, we did not include PC clocks in the present analysis.

We also did not extend the analysis to include protein biomarkers, such as Gadd EpiScores [36]. These scores provide useful biological information by predicting circulating protein levels and disease risk, but their target quantity differs from biological age. The objective of this study was the cross-platform calibration of the biological ages and pace of aging calculated using representative epigenetic clocks. Our selection of Horvath, Hannum, PhenoAge, GrimAge, GrimAge v2, and DunedinPACE was intended to cover widely used first-, second-, and third-generation epigenetic clocks.

This study had certain limitations. First, the samples used to develop and validate the calibration models were obtained from a Japanese population, and additional external validation is required to ensure generalizability to other ethnic populations. Second, we targeted only blood samples, and the utility of these calibrations in other tissues remains unclear. Third, although this study focused on healthy adults, different patterns may exist in disease populations or in older cohorts. Fourth, since coverage of the 20, 000-CpG DunedinPACE preprocessing set differed between EPICv2 (86.9%) and MSA (48.4%), the DunedinPACE results should be interpreted in light of the model-specific imputation and preprocessing implemented in Biolearn.

Furthermore, the EPICv2 and MSA measurements were generated at different facilities in both the training and validation datasets. Therefore, part of the observed cross-platform differences may reflect institution-specific batch effects. Nevertheless, the direction and magnitude of the discrepancies were similar in the training and validation datasets, despite the two datasets being processed at different pairs of institutions. This suggests that the main findings are unlikely to be explained solely by institution-specific batch effects, although complete separation of the platform and batch effects was not possible.

Future research requires large-scale validation, including more diverse populations and tissue types. Calibration models that incorporate more clinical variables and a detailed analysis of the technical factors that cause cross-platform differences are required. Furthermore, for newly developed epigenetic clocks, it is necessary to continuously evaluate cross-platform comparability and update the calibration methods as needed.

In conclusion, this study reveals systematic differences between the EPICv2 and MSA arrays and provides practical calibration methods based on linear regression models, mapping MSA-based epigenetic ages and the pace of aging to EPICv2-based values. Our validation results suggest that calibration is useful for Horvath, Hannum, PhenoAge, GrimAge, and DunedinPACE, whereas routine calibration is not supported for GrimAge v2. The developed calibration models are expected to advance epigenetic research by enabling comparisons of epigenetic ages measured across multiple platforms.

## Acknowledgements

We extend our deepest gratitude to Sawako Hibino, whose efforts in coordinating and collecting blood samples formed the basis of the dataset used in this study. We are profoundly thankful to Genki Yoshikawa, who processed the raw DNA methylation array data and generated the *β*-values necessary for conducting the analyses presented in this study. The contributions of these individuals were indispensable for the successful execution of this study. We would like to thank Editage (www.editage.jp) for the English language editing.

## Code Availability

The Dockerfile and the full pipeline scripts are available from the GitHub repository: https://github.com/tatsumashoji/methylation-beta-pipeline/tree/paper_msa_epicv2_grimagev1. The R scripts used to analyze the data are available from the GitHub repository: https://github.com/t-yui/correctEpiAge-MSA2EPICv2.

## Data Availability

Individual-level epigenetic age data will be made available upon reasonable request.

## CRediT Authorship Contribution Statement

**Yui Tomo**: Conceptualization, Methodology, Software, Investigation, Writing – original draft. **Tatsuma Shoji**: Formal analysis, Software, Investigation, Data curation, Writing – review & editing. **Ryo Nakaki**: Conceptualization, Investigation, Data curation, Writing - review & editing, Project administration, Supervision. All authors have approved the manuscript.

## Declaration of the Use of Generative AI and AI-assisted Technologies

The authors used ChatGPT, Claude, and Gemini to assist with editing and language during the preparation of this manuscript. The manuscript was edited by the authors using these AI tools. The authors take full responsibility for the content of this manuscript.

